# Impact of IgG isotype on the induction of antibody-dependent cellular phagocytosis of HIV by human milk leukocytes

**DOI:** 10.1101/2021.10.14.464395

**Authors:** Alisa Fox, Xiaomei Liu, Susan Zolla-Pazner, Rebecca L. Powell

## Abstract

Approximately 100,000 mother-to-child transmission (MTCT) events of HIV via human milk feeding occur each year (1). However, only about 15% of infants milk-fed by untreated HIV+ mothers become infected, suggesting a protective effect of the milk itself (1, 2). Infants ingest 10^5^-10^8^ maternal leukocytes daily via milk, which remain functional beyond ingestion (3–9). Such function may be elicited by maternal milk antibody (Ab). Though IgA is dominant in milk, most HIV-specific milk Abs are of the IgG subclass, highlighting the importance of investigating the function of each IgG isotype in the milk context (10–16). Though Ab effector function mediated by the constant (Fc) domain via interaction with Fc Receptors (FcRs), such as Ab-dependent cellular phagocytosis (ADCP), are critical in protecting against HIV infection, ADCP is largely unexplored as it relates to mitigation of MTCT (17–21). Presently we report the ADCP activity of milk leukocytes against HIV particles and immune complexes (ICs), using 57 unique samples from 34 women, elicited by IgG1/2/3/4 of monoclonal (m)Ab 246-D. Granulocyte ADCP of HIV was most potent compared to other phagocytes when elicited by IgG1/3/4. IgG1/3 activated granulocytes similarly, exhibiting 1.6x-4.4x greater activity compared to IgG2/4, and a preference for virus compared to ICs. Notably, CD16-monocyte ADCP of a given target were unaffected by isotype, and CD16+ monocytes were poorly stimulated by IgG1. IgG2/4 elicited potent IC ADCP, and in terms of total leukocyte IC ADCP, IgG4 and IgG3 exhibited similar function, with IgG4 eliciting 1.6x-2.1x greater activity compared to IgG1/IgG2, and CD16+ monocytes most stimulated by IgG2. These data contribute to a more comprehensive understanding of Fc-mediated functionality of milk leukocytes, which is critical in order to develop therapeutic approaches to eliminating this route of MTCT, including mucosal administration of mAbs and/or a maternal vaccination aimed to elicit a potent milk Ab response.

## Background

It is evident that although human milk is a vehicle for Mother-to-child transmission (MTCT) of HIV, its virus-killing properties are significant, and unless access to clean water and appropriate infant formula is reliable, the World Health Organization does not recommend cessation of breastfeeding for HIV-infected mothers (1, 2). Milk-derived cytokines, defensins, mucins, erythropoietin, long-chain polyunsaturated fatty acids, oligosaccharides, and lactoferrin have been found to inhibit HIV in vitro, with some being directly correlated with reduced transmission in vivo. Importantly, HIV-specific antibodies (Abs) in milk have been correlated with reduced MTCT, but what remains largely unexplored is the contribution of the cellular fraction of human milk (13).

Human milk is comprised of maternal cells that are >90% viable (22). Cell composition is impacted strongly by stage of lactation, health status of mother and infant, and individual variation, which remains poorly understood (22–25). Given that human milk contains ~10^2^–10^5^ leukocytes/mL, it can be estimated that breastfed infants ingest ~10^5^–10^8^ maternal leukocytes daily (7). Various *in vivo* studies have demonstrated that maternal leukocytes provide critical immunity to the infant and are functional well beyond these sites of initial ingestion (3–9). All maternally-derived cells ingested by the infant have the potential to perform immune functions alongside or to compensate for the infant’s own leukocytes (26). This is likely in conjunction with other components, such as maternal Abs.

Many Abs that are unable to prevent viral entry *per se* are capable of binding to the viral surface or to viral proteins on the surface of infected cells to facilitate a variety of anti-viral activities that ultimately contribute to viral clearance. These functions are mediated by the ‘constant’ region of the immunoglobulin molecule – the crystallizable fragment (Fc) – via interactions with Fc receptors (FcRs) found on virtually all innate immune cells (27). Human milk contains all classes of immunoglobulins, with IgA being dominant (~90% of total Ab); however, most studies have found all HIV-specific Ab in the milk of lactating HIV-infected women to be IgG, a finding which extends to non-human primates (NHPs) and highlights the importance of investigating the function of the various IgG isotypes in milk (10–16). Fc-mediated activity of human milk IgG has been associated with reduced risk of MTCT of HIV, and follow-up analyses of RV144 - the only clinical HIV vaccine trial to show significant efficacy, and various other immune correlate analyses have demonstrated that Fc-mediated Ab effector functions such as Ab-dependent cellular phagocytosis (ADCP) are critical in protecting against HIV infection (17–21). Importantly, FcR allelic variants, particularly for FcγRIIa (implicated in IgG-mediated ADCP against HIV-1), have been shown to be related to disease progression, risk of infection, and/or MTCT of HIV, and it has been shown that IgG from HIV-infected controllers exhibits superior ADCP activity compared to IgG from viremic non-controllers, further highlighting the likely important role of ADCP in mitigating HIV replication (28, 29).

Neutrophils, monocytes, macrophages, and dendritic cells (DCs) are the “professional” phagocytes; these cells have been identified or are purported to be found in human milk (22, 30). We have undertaken a study to assess the ADCP capacity of human milk leukocytes against HIV. Our previous pilot study confirmed milk leukocytes as capable of ADCP of HIV Env gp120-coated microbeads, and demonstrated that most activity was granulocyte-driven (31, 32). In the present study we report our analysis of the ADCP activity of milk leukocytes in 57 unique samples from 34 women, elicited by recombinant IgG1/2/3/4 of monoclonal (m)Ab 246-D, a weakly-neutralizing Env gp41 (cluster 1)-specific Ab - the type of Ab commonly elicited by natural HIV infection.

## Materials and Methods

### Participant information

Participants were recruited using social media and informed consent was obtained in accordance with the ethical and institutional board approval under the guidance and authorization of Mount Sinai’s Program for the Protection of Human Subjects (PPHS) using an IRB-approved protocol for obtaining human milk samples (STUDY-17-01089). Individuals were eligible to have their milk samples included in this analysis if they were lactating, feeling well, and living in New York City. Participants were excluded if they had any acute or chronic health conditions affecting the immune system. Milk was expressed using manual or electric pumps in participants’ homes just prior to sample pickup using clean containers. Milk was used for ADCP experiments within 4 hours of expression.

### Recombinant Ab production

mAbs 246-D and 3865 were originally isolated from human peripheral blood mononuclear cells (33, 34), and were expressed recombinantly in 293T cells for this study as human IgG1, IgG2, IgG3, IgG4 using IgG plasmids (Invivogen) by conventional cloning techniques as previously described (35, 36).

### *GFP-HIV* target virus

The infectious molecular clone (HIV-GFP_11056_) used for these experiments carries a Gag-interdomain green fluorescent protein (GFP), and these GFP viruses were originally described by Dr. Benjamin Chen who generously supplied the virus in the present study (37). Virus was produced by standard transfection of 293T cells, sucrose-gradient purified, and quantified by p24 ELISA as previously described (37).

### ADCP assay

This primary milk cell assay was adapted from Ackerman et al (38) and has been described previously using latex beads (32). Milk samples were centrifuged at 600*g* for 15 min, cell pellet was washed 2x with Hank’s balanced salt solution (HBSS) without calcium and magnesium with centrifugation at 350g for 10 min. Care was taken to gently resuspend cell pellets to avoid granulocyte activation and apoptosis. For virus ADCP, 50ul of 300ng/mL (by p24 quantification) GFP-HIV was aliquoted in 96-well plates. Titrated (200ug/ml – 12.5ug/ml) mAb in 50ul was incubated with virus for 2hrs at 37°C. Fifty thousand milk cells were added per well and plate was incubated as above. For IC ADCP, 100ug/mL anti-human IgG(Fc) (Life Technologies) was added to virus-Ab complexes and incubated for 30 min at 37°C prior to the addition of cells as described in (39). Control experiments to distinguish between virions bound to the cell-surface and those phagocytosed by cells were performed by pre-incubating milk cells with 10ug/mL of the actin inhibitor Cytochalasin-D (Sigma) and/or 50ug/mL of FcR blocking agent FcBlock (BD). After incubation, cells were washed and incubated in Accutase cell detachment solution for 10 min at room temperature to gently remove any surface-bound virus while preserving epitopes for flow cytometry analysis (Innovative Cell Technologies; as described in (39)). Cells were then washed and stained. For all CD45+ leukocytes, as well as for each of the phagocyte subsets, ADCP activity was measured as the percent of GFP+ cells of that cell type. ADCP scores for total CD45+ leukocytes were calculated as [(MFI of GFP+ cells) x (% GFP+ cells of the CD45+ population)], and for each phagocyte subset were calculated as [(MFI of GFP+ cells) x (% of total CD45+ cells in the GFP+ population)], in order to provide a relative assessment of each cell population’s overall phagocytic contribution within each unique sample. Experiments for each milk sample were performed in duplicate. Scores at each mAb dilution were plotted using GraphPad Prism and Area under the Curve (AUC) values were calculated for each experiment. AUC scores were normalized by subtracting those obtained for each cell type using the irrelevant anthrax mAb 3865 of the same class.

### Gating strategy

Analysis was performed using an LSR Fortessa. Cells were stained as directed by the manufacturer for CD45, CD3, CD14, CD16, CD20, CD56, and MHC-II (HLA-DR, DP, DQ) (BD) (30, 40) (41). Initial gating and viability dye (BD) was used to exclude doublets and dead cells. A side scatter (SSC) vs. CD45 plot was then used to identify CD45+ leukocytes. FSC/SSC was used to identify monocytes followed by gating to exclude CD3+ T cells, CD20+ B cells, and CD56+ NK cells. MHC-II/CD14/CD16 staining was employed to differentiate classical monocytes (CD14+CD16−) from activated monocytes/macrophages (CD14+CD16+ and CD14dimCD16+) (41, 42). DCs were identified by selecting from the FSC/SSC plot to include both lymphocytes and monocytes, followed by gating on MHC-II+ cells after exclusion of CD3+ T lymphocytes, CD14+ monocytes, CD20+ B lymphocytes, and CD56+ NK cells. Granulocytes were identified based on the FSC/SSC plot and further confirmed by exclusion of other cell markers as above, and positive CD16 stain. Gating strategy is shown in Fig. 1.

**Figure 1:**
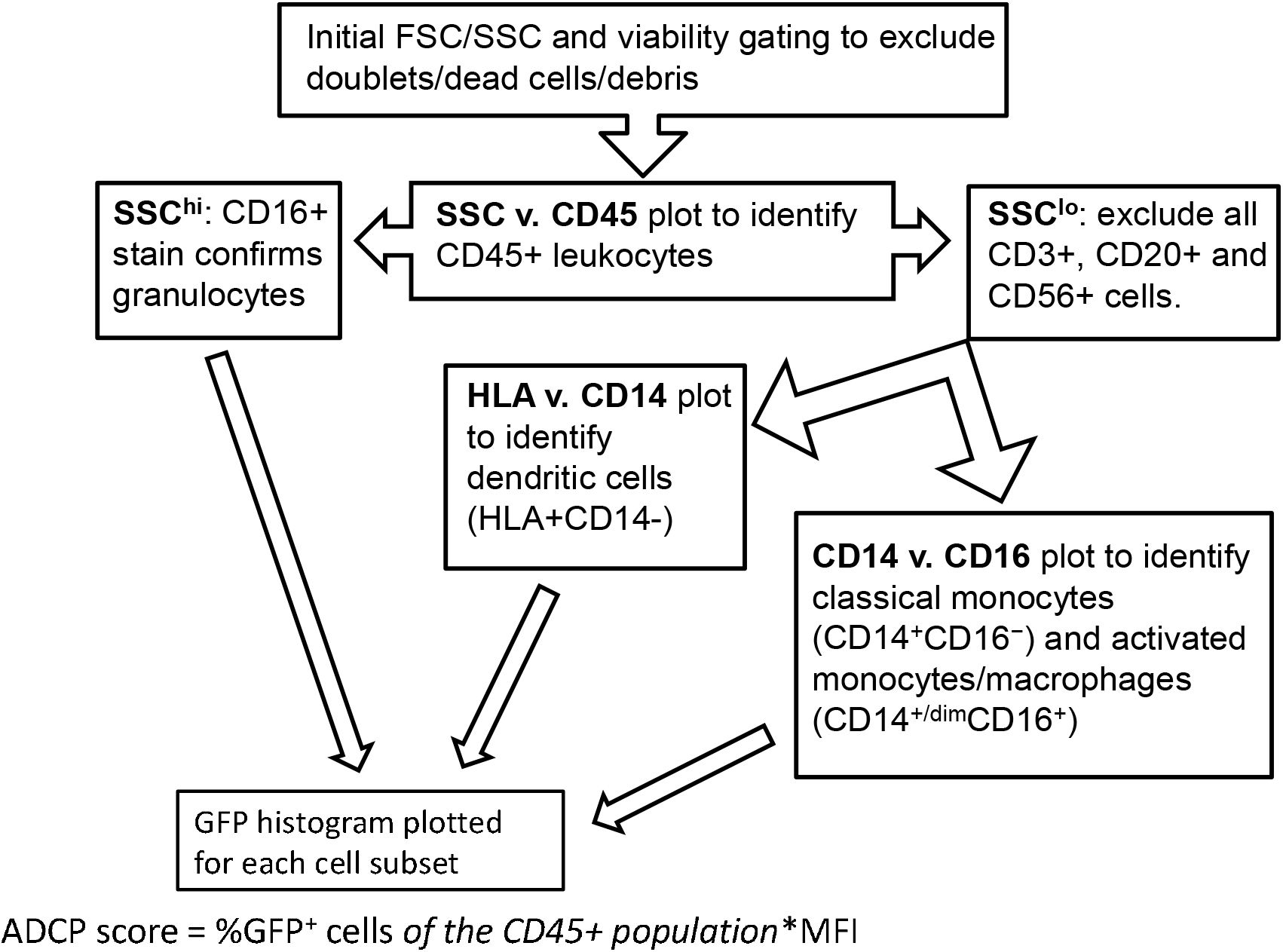
Gating strategy. Flow cytometery was performed using an LSR Fortessa (BD) with initial gating on FACSDiva software. Analytical gating was performed using FCS Express.

## Results

### Sample data

Fifty-seven unique milk samples were obtained from 34 different women who were between 1 – 24 months postpartum at the time of milk collection (mean, 8.1 months; Fig. 2a), yielding 42mL – 250mL of milk per sample (mean, 88.4mL; Fig. 2b). Samples contained between 12,054 and 707,143 total cells/mL of milk (mean, 69,291 cells/mL; Fig. 2c), which included 0.02% - 33% CD45+ leukocytes (mean, 6.83%; Fig. 2d). On average, classical monocytes (SSC^int^CD45^+^CD3^−^CD20^−^CD56^−^CD16^−^CD14^+^) and granulocytes (SSC^hi^CD45^+^CD16^+^) accounted for similar proportions of the CD45+ leukocytes (15.1% and 11.8%, respectively), while the proportion of activated monocytes/macrophages (SSC^int^CD45^+^CD3^−^CD20^−^CD56^−^CD16^+^CD14^+/dim^) was significantly lower compared to the classical monocytes (mean, 6.03%), and the proportion of dendritic cells (DCs; SSC^lo/int^CD45^+^CD3^−^CD20^−^CD56^−^CD16^+/−^D14^−^HLA^+^) was lowest of all subsets identified – significantly lower than both classical monocytes and granulocytes (mean, 3.18%; Fig. 2e).

**Figure 2:**
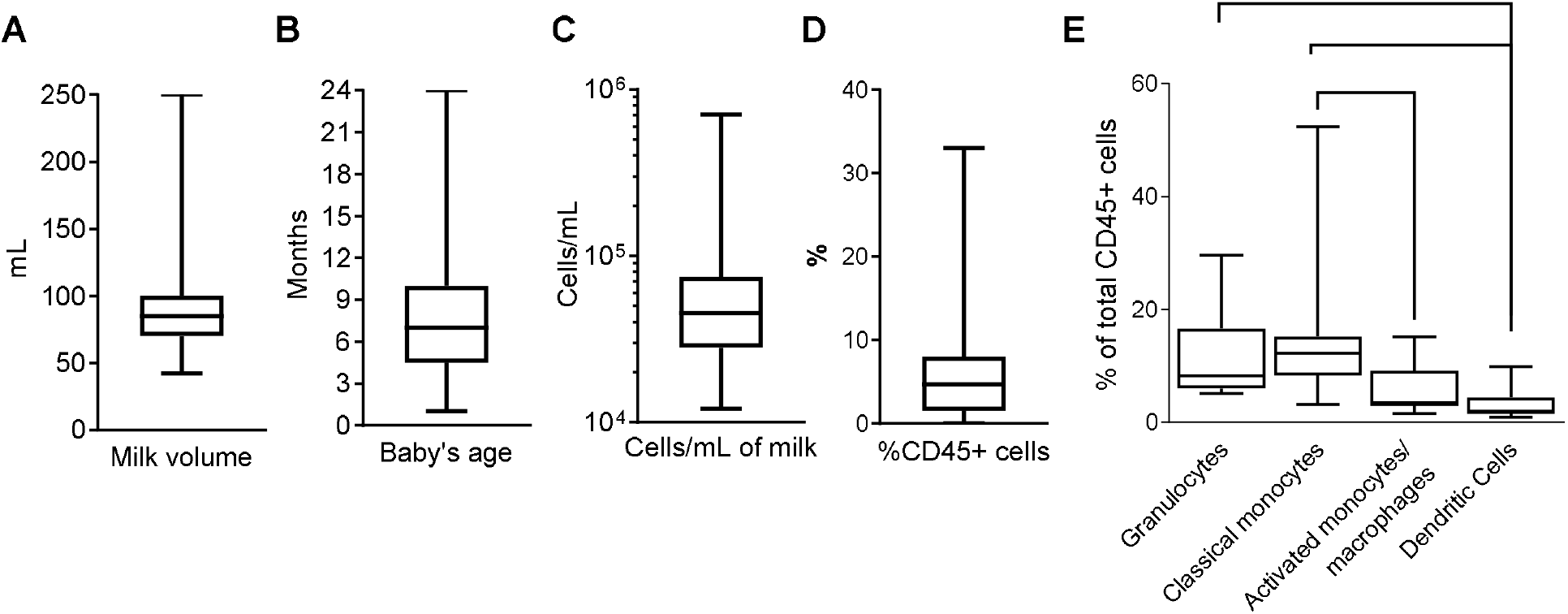
Sample data. (a) Milk volume collected for each sample. Milk was expressed just prior to sample pickup and used within 4h. (b) Baby’s age/time post-partum of study participant. (c) Total cell concentration for each sample. Milk was centrifuged at 600*g* for 15 min, and cell pellet washed with HBSS at 350g. Cells were counted, and stained as directed (BD). (d) Percent CD45+ cells of total cell population as determined by flow cytometry. (e) Phagocyte subset populations expressed as percent of total CD45+ cells. CD45+ cells and phagocyte subsets were identified as described in Fig. 1. Box and whisker plots with minimum, maximum, and mean values are shown. Two-tailed Mann-Whitney tests were used to compare phagocyte populations. ***0.0005≥p>0.0001. *p=0.028.

### ADCP activity of 246-D-opsonized virus and ICs as compared by effector cell type

Total CD45+ leukocytes as well as each phagocyte subset were assessed for their ADCP capacity against the GFP-HIV target virus opsonized with titrated mAb 246-D. Blocking experiments with CytochalasinD and FcBlock demonstrated that ADCP activity was actin and FcR dependent, with residual activity below that of the 3865 mAb control (Fig 3). ADCP was performed using milk cells isolated from 11-19 unique samples for each IgG isotype tested.

**Figure 3:**
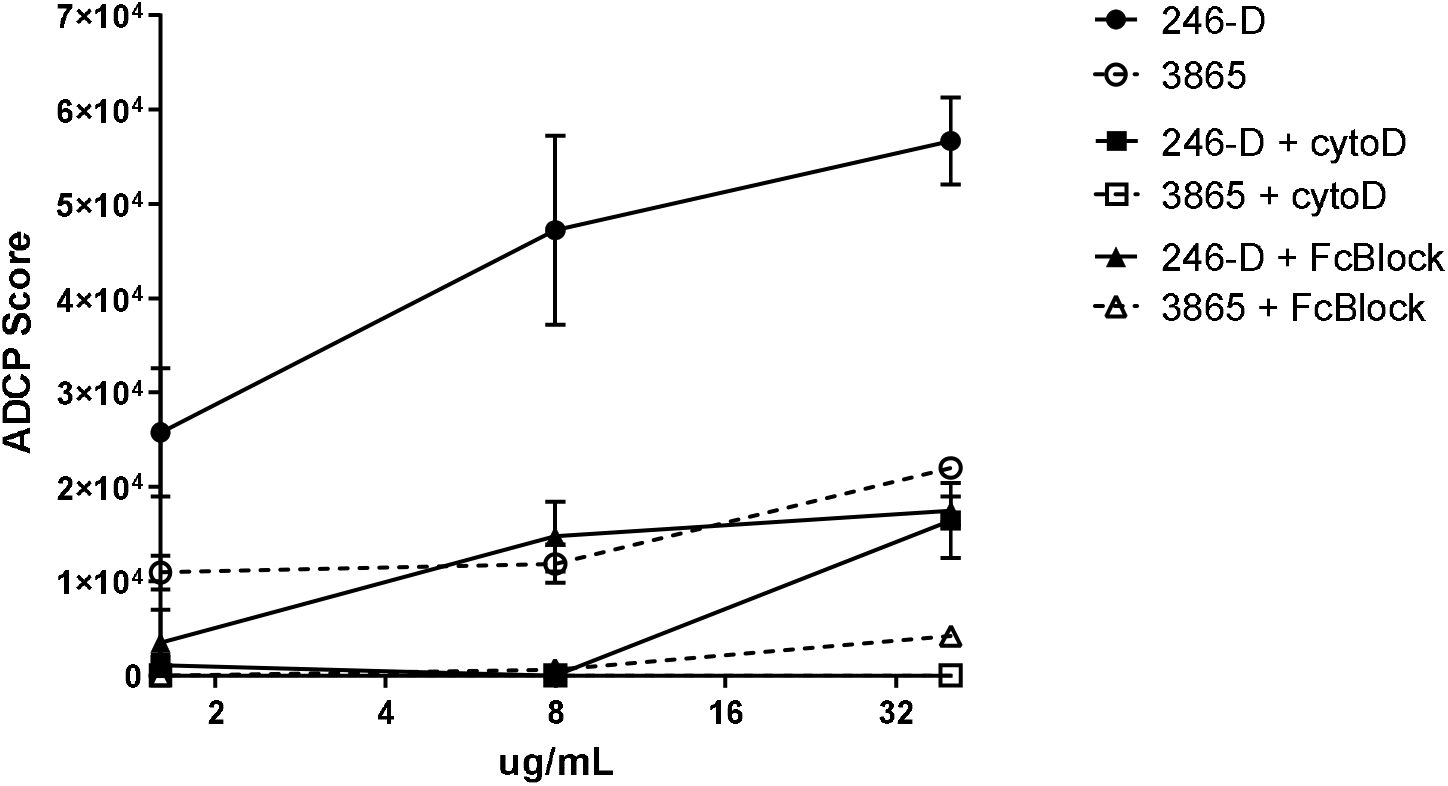
ADCP activity measured using milk phagocytes is actin and FcR dependent. 300ng/mL GFP-HIV per well was incubated with titrated IgG1 mAb for 2hrs at 37°C. Meanwhile, 50,000 milk cells per well in a separate plate were prepared, with certain wells pre-incubated with 10ug/mL of the actin inhibitor Cytochalasin-D (Sigma) or 50ug/mL of FcR blocking agent FcBlock (BD) for 30min at 37°C. Cell plates were washed and incubated with virus/mAb complexes. Plates were washed and incubated in Accutase cell detachment solution, then washed and stained. ADCP activity was measured as the percent of GFP+ cells, with ADCP scores calculated as [(MFI of GFP-positive cells) x (% of GFP+ cells of CD45+ population)]. Data shown was measured for CD45+ population. Mean values of duplicate experiments are shown with SEM.

#### IgG1-elicited ADCP

As IgG1, 246D-elicited granulocyte ADCP of HIV-GFP was the most dominant, and this activity was significantly greater than that of all other phagocyte subsets (6.7x – 57x greater activity; p<0.0001; Fig. 4a). Activated and classical monocyte ADCP were of similar potencies, wherein both monocyte types exhibited significantly more potent virus ADCP activity compared to DCs (6.7x – 8.4x greater activity; p=0.001 and p=0.009, respectively; Fig. 4a). Similarly, granulocyte ADCP of the larger anti-Fc-complexed HIV-GFP immune complexes (ICs) was the most potent (1.7x – 13x greater activity; p=0.005 and p<0.0001, respectively), although this activity was not significantly greater than that of the activated monocytes in this case (Fig. 4a). In fact, against these larger targets, activated monocyte activity was significantly increased compared to that of the classical monocytes and the DCs (3.2x – 7.6x greater activity; p=0.023 and 0.0004, respectively), while classical monocyte and DC activities were similar (Fig. 4a). When comparing ADCP potency against opsonized virus vs. IC for each cell type, only granulocytes exhibited a significant preference, for the smaller, ‘free’ virus (1.6x greater activity; p=0.024).

**Figure 4:**
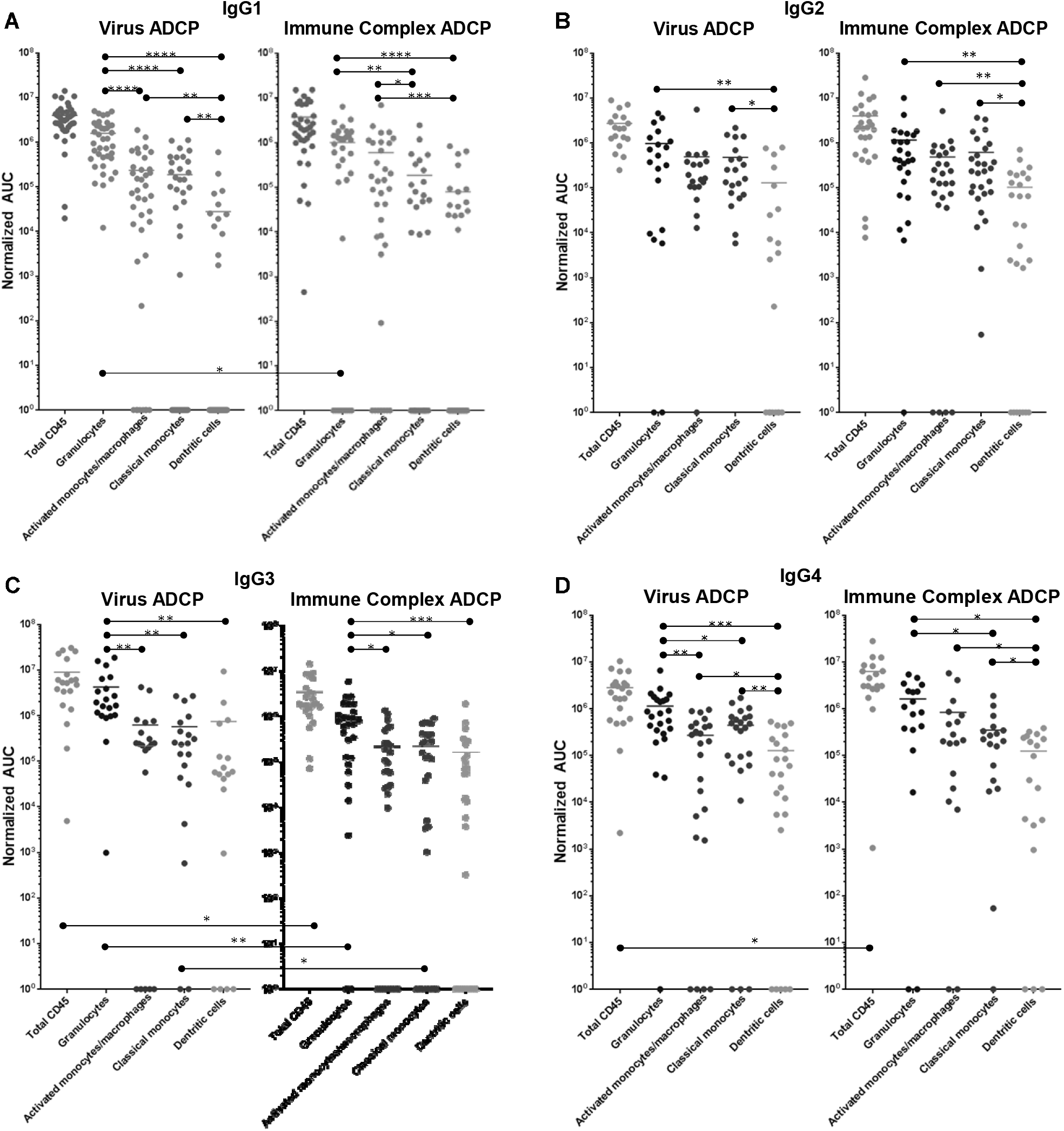
ADCP activity of 246-D-opsonized virus and ICs as compared by effector cell type. (a) IgG1 data. (b) IgG2 data. (c) IgG3 data. (d) IgG4 data. ADCP was performed as described above. ADCP scores at each mAb dilution were plotted using GraphPad Prism and Area under the Curve (AUC) values were calculated for each experiment. AUC scores were normalized by subtracting those obtained for each cell type using the irrelevant anthrax mAb 3865 of the same class. Each data point represents cells from a unique milk sample. Mean AUC are shown. Two-tailed Mann-Whitney tests were used to compare data. ****p<0.0001; ***0.0009>p>0.0001. **0.009>p>0.001;* 0.05>p>0.01.

#### IgG2-elicited ADCP

As IgG2, 246D elicited granulocyte ADCP of HIV-GFP significantly more potently than DCs (7.5x greater activity; p=0.0076; Fig. 4b), but not compared to any other cell type. Activated and classical monocyte ADCP were of similar potencies, though classical monocytes also exhibited significantly more potent virus ADCP activity compared to DCs (3.7x greater activity; p=0.0125; Fig. 4b). Granulocyte ADCP of the HIV ICs was again only significantly greater than that of the DCs (11.1x greater activity; p=0.0058; Fig. 4b). Classical and activated monocyte ADCP was similar, while both cell subsets exhibited more potent ADCP than DCs (4.7 – 6x greater activity; p=0.0046 – 0.029; Fig. 4b). When comparing ADCP potency against virus vs. IC for each cell type, no significant differences were found.

#### IgG3-elicited ADCP

As IgG3, 246D elicited significantly greater granulocyte ADCP of the virus particles compared to all other cell types (5.7x – 6.8x greater activity; p=0.0023 – 0.014; Fig. 4c). All other phagocyte subsets exhibited similar activities. Granulocytes also exhibited significantly greater ADCP compared to all other cell types against the HIV-ICs when elicited by IgG3 (4.1x – 5.9x greater activity; p=0.0006 – 0.011; Fig. 4c). As observed for the virus targets, all other phagocyte subsets exhibited similar activities. Notably, total CD45+, granulocyte, and classical monocyte ADCP activity was significantly greater against the virus targets compared to the ICs (2.5x – 4.4x greater activity; p=0.0096 – 0.0225).

#### IgG4-elicited ADCP

As IgG4, 246D elicited significantly greater granulocyte ADCP compared to all other cell types (2.6x – 9x greater activity; p=0.0008 – 0.022; Fig. 4d). Additionally, both monocyte subsets exhibited greater ADCP activity compared to the DCs (2.1x – 3.5x greater activity; p=0.001 – 0.026; Fig. 4d). Against the HIV-ICs, granulocytes exhibited significantly greater ADCP than classical monocytes and DCs (4.6x – 13x greater activity; p=0.0003 – 0.0019; Fig. 4d), while both monocyte subsets exhibited greater activity compared to DCs (2.8x – 6.7x greater activity; p=0.031 – 0.032; Fig. 4d). Total CD45+ ADCP activity was significantly greater against the ICs when elicited by IgG4 compared to the virus targets (2.4x greater activity; p=0.0096 – 0.0225).

### ADCP activity as compared by IgG isotype

#### Virus ADCP

When including all CD45+ leukocytes, it was evident that 246-D as IgG3 elicited ADCP of virions significantly better than as IgG2 or IgG4 (3.2x – 3.3x greater activity; p=0.017 – 0.026; Fig. 5a). In terms of granulocyte ADCP, IgG1 activity was significantly greater than IgG2 activity (1.6x greater activity, p=0.030), while similar to total CD45+ leukocytes, IgG3 elicited granulocyte virion ADCP more strongly than IgG2 or IgG4 (3.7x – 4.4x greater activity; p=0.0013 – 0.0029; Fig. 5a). No significant differences in ADCP activity was found for activated monocytes/macrophages when comparing IgG isotypes (Fig. 8a). In contrast, classical monocyte ADCP was elicited more strongly by IgG2, IgG3, and IgG4 compared to IgG1 (2.4x – 3.1x greater activity, p=0.0051 – 0.0297; Fig. 5a). Similarly, dendritic cell ADCP was also elicited more strongly by IgG2, IgG3, and IgG4 compared to IgG1 (4.6x - 26.7x greater activity; p=0.0336 - p<0.0001; Fig. 5a).

**Figure 5:**
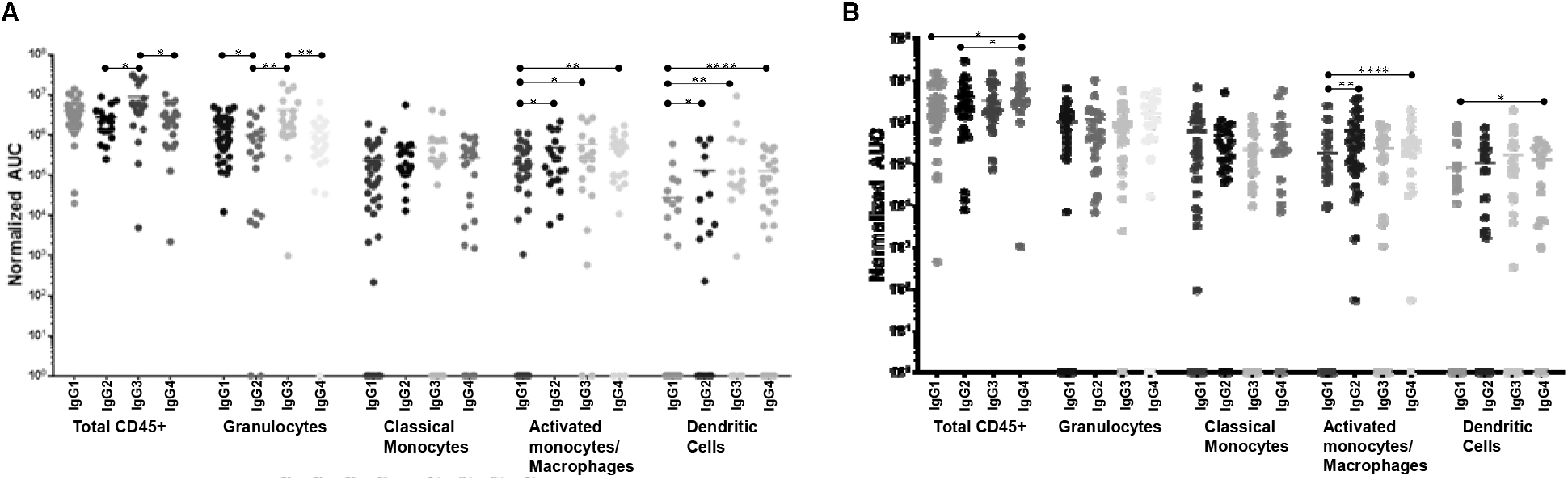
ADCP activity of each phagocyte subset as compared by IgG isotype. (a) Virus ADCP. (b) Immune complex (IC) ADCP. Experiments and analyses were performed as in Fig. 4.

#### IC ADCP

Measuring total CD45+ cells, IgG4 was the most potent at eliciting IC ADCP compared to IgG1 and IgG2 (1.6x – 2.1x greater activity; p=0.0176 – 0.0399; Fig. 5b). Granulocytes and activated monocytes/macrophages exhibited no significant differences in ADCP activity when comparing the IgG isotypes. Classical monocyte IC ADCP was elicited more strongly by IgG4 and IgG2 compared to IgG1 (1.9x and 3.4x greater activity; p=0.0035 and p<0.0001, respectively; Fig. 5b). Lastly, it was also apparent that IgG4 elicited more potent IC ADCP by dendritic cells compared to IgG1 (1.6x greater activity; p=0.0121; Fig. 5b).

## Discussion

Each phagocyte subset expresses a unique FcγR repertoire that includes FcγRI, FcγRIIa/b/c, and/or FcγRIIIa/b (43). As well, FcR expression and function of phagocytes can vary by their tissue localization; for example, HIV-1-specific ADCP activity of colon-resident macrophages is virtually non-existent compared to those found in the cervix (44). Most FcγRs have a measurable affinity for all IgG isotypes, though selectivity for a given isotype certainly exists; moreover, most FcRs mediate cytoplasmic signaling, including FcγRIIa/FcγRIIc, which possess immunoreceptor tyrosine-based activation motif (ITAM) domains, and FcγRI/FcγRIIIa that signal via a γ2 chain containing an ITAM domain (18, 45). Binding to most FcγRs requires the Ab to be at least dimerized, i.e., part of an immune complex, and in this form, affinity experiments have shown that both IgG1 and IgG3 bind all FcγRs comparably, with only a slightly better binding profile exhibited by IgG3 (45). Even FcγRI, which can bind IgG monomers with high affinity, requires receptor clustering and cross-linking to signal effectively (46).

In the present study, granulocytes were the most potently induced phagocyte against the virus target for most IgG isotypes, consistent with our previous findings using bead-based targets (32). IgG2 did not appear to elicit granulocyte ADCP more potently than monocyte ADCP, possibly due to the apparent lack of binding affinity of IgG2 for FcγRIIIb, which is dominantly expressed on neutrophils, and for FcγRI, which neutrophils express when overstimulated (45, 47). Granulocyte dominance was less apparent for the IC targets, likely due to the extensive crosslinking of Fc-FcR interactions leading to similar levels of ADCP across the cell types tested, with the exception of IgG3, which induced highly potent ADCP by granulocytes of ICs compared to other cell types. Otherwise, IgG1/2/4 in this case more similarly stimulated granulocytes and activated monocytes/macrophages, or even classical monocytes in the case of IgG2. Overall, dendritic cells exhibited the lowest relative activity in terms of both virus and IC targets, though this would be expected given these cells comprise the smallest fraction of phagocytes analyzed.

The ADCP scores presented herein are a relative assessment of each cell population’s overall phagocytic contribution within each unique sample, as the scores take into account the percent of total CD45+ leukocytes each phagocyte subset comprises. The ADCP activity measured here is therefore a biologically applicable indication of the most relevant cells and isotypes to target for further studies that make use of these data. However, when directly comparing the ADCP activity induced by each IgG isotype for a given cell type, the absolute phagocytic potency can be better examined. In this case, it was apparent that examining all CD45+ leukocytes together, IgG3 and IgG1 elicited similar activity against the virus target, which is likely a reflection of the potent activity of granulocytes whose dominant FcγRIIIb expression limited IgG2 and IgG4-elicited ADCP activity due to the low affinity of this interaction even when measured in cell-surface, cross-linked binding studies (45). Classical CD14+CD16-milk monocytes were the least impacted by isotype, suggesting the high levels of Ab used in these experiments to opsonize the virus targets facilitated cross-liking of FcγRIIa to such an extent that differential affinities for each isotype was essentially irrelevant. Indeed, even moderate crosslinking of Ab acting on FcγRIIa has been shown to minimize any binding differences between the ‘high affinity’ IgG1/IgG3 and ‘low affinity’ IgG2/IgG4 (45). Though primary blood monocytes express high levels of FcγRI, which exhibits very poor IgG2 binding even when cross-linked, these data suggest that FcγRIIa activity accounted for this difference, and/or that milk monocyte FcγRI expression may be relatively low (48). IgG isotype also seemed to have minimal impact on activated monocyte/macrophage and DC ADCP; however, these cell types were most poorly stimulated by IgG1. Reasons for this are not clear, though it is possible that combined with the very limited expression of FcγRI expression on these cells (particularly DCs), IgG1 induction of ADCP may have exhausted the FcγRs and/or overstimulated the inhibitory FcγRIIb. Notably, *in vivo,* the high-affinity FcγRI may be consistently bound by monomeric IgG; as such, the lower affinity FcγRs are responsible for most observed Fc-mediated activities (49).

Overall, IC ADCP appeared to be best mediated by IgG4. This was most apparent for DCs, as well as for activated monocytes/macrophages, which also exhibited potent IgG2 activity. However, granulocyte activity against ICs appeared to be unaffected by IgG isotype. Granulocytes exhibited more potent ADCP of the smaller virus targets when elicited by the high affinity IgG1 and IgG3, but no difference in terms of IgG2 or IgG4, again likely a reflection of FcγRIIIb binding preferences. The significant IC crosslinking and/or target size appeared to overcome the typical lack of FcγRIIIb-IgG2/4 affinity, though it may be that the observed activity was triggered strongly by granulocyte FcγRIIa despite the lower expression levels expected on these cells.

It was apparent from these data that with extensive crosslinking of larger ICs, isotype of the effector IgG becomes considerably less relevant in the context of gp41-mediated ADCP. In fact, isotypes less classically associated with effector function, particularly IgG4, appear to function well in this context. Given the overall low numbers of DCs, they likely do not have a major impact on ADCP activity in milk, though they certainly can perform this function and likely contribute to viral clearance. As also observed in our previous study using gp120-coated bead targets, granulocytes are potent mediators of ADCP in milk, and the present data demonstrates that these cells should be targeted therapeutically to mitigate MTCT of HIV via breastfeeding. Depending on the therapeutic approach, be it mucosal administration of mAbs to elicit powerful enhancement of the natural passive immunity of HIV+ mothers’ milk, or a therapeutic maternal vaccine aimed to elicit a potent milk Ab response, it is evident that granulocyte preferences, in particular, for IgG isotype in terms of elicitation of ADCP should be considered. Should IgG3 be used therapeutically, these data suggest that granulocytes will be potently activated, though very high viral loads and/or very high mAb doses that might generate large ICs might reduce the potency of this effect, and careful titration experiments would have to be done *in vivo*, also taking into account the short half-life of this isotype (27). These data also suggest IgG1 may not be an ideal choice for such therapeutics to fully harness the ADCP potential of milk monocytes, while high doses of IgG4 in a high viral load context may induce ADCP by all milk phagocytes. As IgG4 induction tends to occur after repeated and prolonged immunogen exposure, its ADCP functionality may become relevant should frequent maternal vaccine boosters be used (50).

Importantly, though most HIV-specific Abs in milk have been found to be IgG, the dominant Ab class in milk is sIgA, and the Fc-mediated function of (s)IgA must be better understood, as it may be relevant therapeutically. As well, the present data has highlighted that the milk leukocyte FcR expression profiles should be further examined in subsequent studies. Given the substantial gap in present knowledge of the potential contribution of ADCP activity by milk phagocytes to prevention of MTCT of HIV, it is critical to develop a multidimensional, comprehensive understanding of ADCP by these relevant primary cells.

## Acknowledgements

We are indebted to our milk donors. HIV-GFP was generously gifted from the Benjamin Chen lab. This work was supported by the NIH/NIAID and the Icahn School of Medicine at Mount Sinai.

